# Emotion Dynamics in Reciprocity: Deciphering the Role of Prosocial Emotions in Social Decision-making

**DOI:** 10.1101/2023.12.18.572094

**Authors:** Jaewon Kim, Su Hyun Bong, Dayoung Yoon, Bumseok Jeong

## Abstract

To date, the relevance of prosocial emotions in social decisions based on reciprocity remains poorly understood. Expected and experienced emotions in interoceptive-social dimension, expected offers, and actual acceptance were measured in 476 participants during an ultimatum game consisting of fair, moderate, and unfair offers. We investigated whether participants adjust social decisions according to prediction errors on prosocial emotions and reciprocity. Participants’ acceptance trajectories were explained by prediction errors in dominance, valence, and reward. Participants were categorized into 4 distinct subgroups based on their patterns of reward expectation, acceptance, and emotional experiences before and after the offer. Furthermore, the relationships between prosocial emotions, social decisions, and reciprocity varied across these subgroups. This study’s measurement and analysis of multidimensional trajectories across four affect dimensions reveal that social decisions are influenced by the responder’s perception of partner’s reciprocity, as well as by the subsequent prediction error of basic and prosocial emotion.

## Introduction

“If someone (I) were in my (their) position, they (I) would behave in a similar manner as I (they) would.” This belief and interaction based on are collectively called reciprocity. Reciprocity incorporates a sense of fairness and balance into social decision-making. Studies on social decision-making have investigated neuroeconomic tasks and reported that human social decisions are influenced by economic profit as well as other factors such as a sense of fairness and reciprocity[1-3].

Social decision-making is frequently investigated using an ultimatum game (UG). In UG, a proposer makes an offer to a responder who decides either to accept or reject the offer. If the responder accepts the offer, both players receive the reward that is split by the proposer. If the responder rejects, none of the players receive the reward; hence, the proposer should act more prosocially.

The reward prediction error (RPE), which is defined as the difference between the expected outcome and the proposer’s actual offer, represents the subjective experience of reciprocity. Studies have reported that the RPE influences decisions during neuroeconomic tasks[4, 5]. Recently, predictive emotions, in the form of the emotion prediction error (EPE), were also reported to predict the choices of UG responders[6]. Participants demand more prosocial interaction by punishing their partners when they receive smaller rewards or feel less pleased and more emotionally aroused than expected. In other words, social decision-making during UG is paired not only with the subjective experience of the partner’s reciprocity but also with the responder’s emotional response to it.

Conventionally, valence and arousal have been incorporated into a two-dimensional affect grid to represent basic emotions[7]. In 2021, Heffner, Son, and FeldmanHall adopted this affect grid to divide emotion PE into two dimensions based on the valence prediction error and arousal prediction error[6]. However, in response to prosocial or unfair offers, responders may experience prosocial[8] (e.g., gratitude, shame, embarrassment) as well as basic[9] (e.g., happiness, sadness, anger) emotions.

Studies on affective representation have proposed other dimensions for these prosocial emotions. One of these is dominance, the degree to which an emotion feels in or out of control to a person[10, 11]. Tangney et al. have used another dimension, i.e., focus, which refers to the degree of an emotion’s relevance to oneself or others, along with valence[8]. Therefore, emotions in response to a proposer’s offer could be better explored using four dimensions, namely, valence, arousal, focus, and dominance.

Playing an iterative neuroeconomic game while simultaneously rating the affect, which is called dynamical affective representation mapping (ARM), has made it possible to trace momentary emotions during social decision-making[6, 12]. However, previous studies have typically focused on aggregate summaries of moment-to-moment emotions. A dynamics approach has been primarily employed to examine emotional dynamics associated with specific, isolated behaviours rather than considering the entire spectrum of an individual’s emotional experience. Since a dynamical ARM output involves time series, a measurement of participant responses using a dynamics approach such as a trajectory analysis[13, 14] can provide novel insights into human emotional experience. To the best of our knowledge, this approach has rarely been used in previous studies.

By adopting a dynamics approach and unsupervised neural network classification algorithm, we aim to investigate trajectory of responders’ predictive emotions and social decision-making during UG. Our main hypotheses include: 1) Decisions during UG will be further explained by adding the responder’s emotion prediction error expressed in four affective dimensions to reward prediction errors across different fairness levels. 2) Individual differences of relationship between predictive emotion, social decision, and reciprocity can be identified with affective representation adopting these social dimensions. We predict that there will be groups that show distinct pattern of social decision-making, as well as related experiences of reward expectation, predictive emotions (Fig. 1c). We also predict that groups will have different relationships between predictive emotion, social decision, and reciprocity.

**Figure 1.**
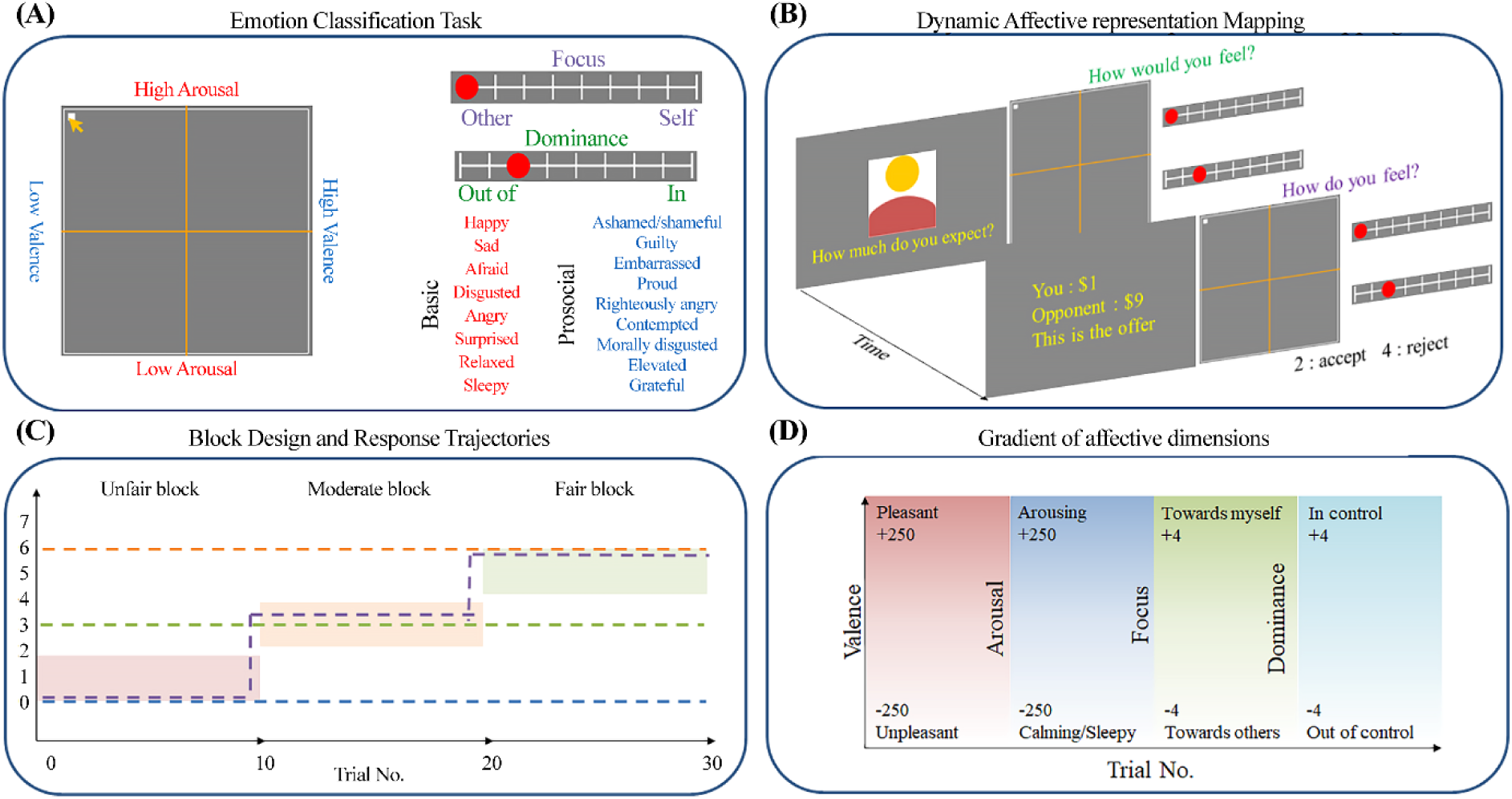
Diagrams of experimental procedures. (A) During the ECT, the participants rate basic and prosocial emotions using valence, arousal, focus, and dominance (affective dimensions). (B) During dynamic ARM with UG, participants are asked to respond with the offer they expect from the proposer (expected reward) and the affective dimensions they would feel if they were given the expected reward (expected emotion). After the offer is extended, the participants rate their affective dimensions at that moment (experienced emotion) and then decide (reward acceptance). (C) Rewards are drawn from pseudorandomised values corresponding to three different fairness levels represented by colored boxes. Examples of potential response trajectories are shown using dashed lines. (D) Conceptual diagram of each affective dimension, along with its gradient.

## Results

### Experimental procedure

The participants played an in-house computerised cognitive task, which included three stages. During the emotion classification task (ECT), the participants rated a series of 18 emotion labels (9 basic emotions, 9 moral emotions) based on dynamical ARM measures (Fig. 1a). These dynamical ARM measures were the valence, arousal, focus, and dominance (affect dimensions). Subsequently, the participants played 30 trials of one-shot UG consisting of three blocks with increasing fairness of offers (Fig. 1b). During each trial, the participants responded to questions regarding (1) how much reward they expect from the proposer as an offer (expected reward) and (2) the affect dimensions they would feel if they were given the expected reward (expected emotion) before the offer. After the offer was made, the participants rated (3) the affect dimensions they actually felt at that moment (experienced emotion) and their (4) UG decision (accept or reject) as a responder (reward acceptance). Details of the experiments are presented in the methods section.

### Unsupervised clustering algorithms reveal common data structure across time series of reward and emotions

Clustering of the participants based on the trajectories of their expected reward (Fig. 2a), reward acceptance (Fig. 2c), and expected (Fig. 2e) and experienced (Fig. 2g) emotions resulted in solutions with equal numbers of components. Clustering for all components resulted in solutions wherein the cluster centroids showed three distinct stationary trajectories with consistently low, middle, and high values and one distinct dynamic trajectory that chased the actual reward (Supplementary Figs. 5 and 6). We labelled each cluster of the expected reward/emotion as pessimistic (PES), fair (FAIR), optimistic (OPT), and reciprocal expectation (RECe) groups, and each cluster of the reward acceptance and experienced emotion as the non-cooperative (NON), indifferent (IND), rational (RAT), and reciprocal experience (RECx) group.

**Figure 2.**
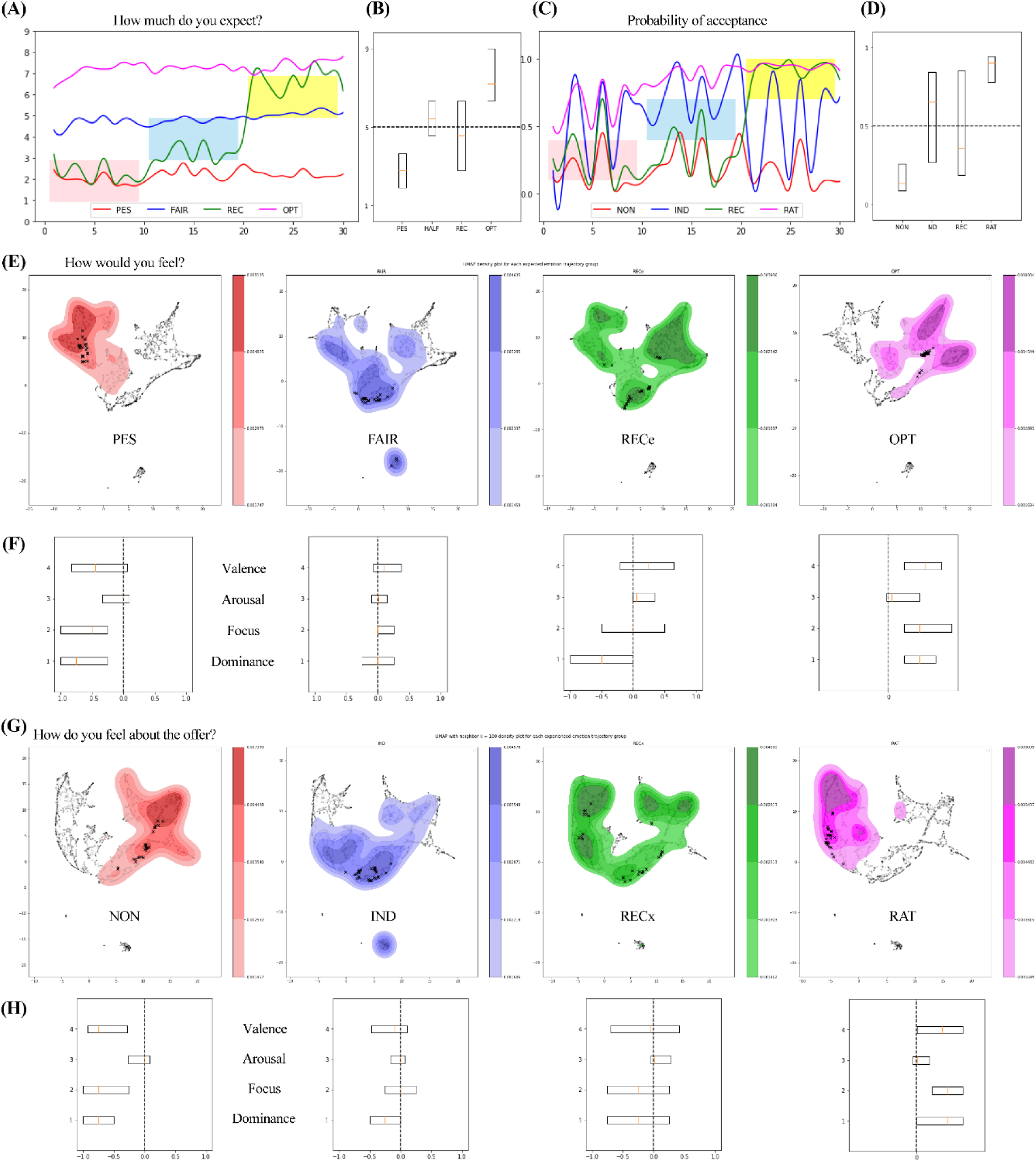
Results of trajectory analyses. (A, C) Trajectories of expected reward and reward acceptance. The X axes represent the trial numbers, and the Y-axes represent the (A) expected offer and (C) probability of acceptance (p_accept) for a given trial. The colored boxes represent unique fairness levels, and the trajectories were smoothed using SciPy for visualisation. (B) Box plot of the expected reward for each group indicated in (A). (D) Box plot of p_accept for each group indicated in (C). (E, G) UMAP embedding of the (E) expected and (G) experienced emotional trajectories of the participants. The black markers indicate the centroid trajectory of each group, and the kernel density estimation plot for each group was overlaid using SciPy (grid size = 600, levels = 5, thresh = 0.3, alpha = 0.8). Each colored line or cloud represents one of the four groups. Groups of the same color across (A)∼(H) display semantically matched trajectories but do not comprise the same participants. (F,H) Box plots of (F) expected and (H) experienced emotions. Abbreviations: PES, pessimistic expectation group; FAIR, fair expectation group; RECe, reciprocal expectation group; OPT, optimistic expectation group; NON, non-cooperative group; IND, indifferent group; RECx, reciprocal experience group; RAT, rational group.

The average expected rewards for each expected reward group were 2.16 (PES, N = 161), 4.85 (FAIR, N = 194), 7.27 (OPT, N = 75), and 4.09 (RECe, N = 46). The mean acceptance ratio for each reward acceptance group were 0.184 (NON, N = 131), 0.579 (IND, N = 105), 0.828 (RAT, N = 163), and 0.489 (RECx, N = 77).

Since the data of the expected and experienced emotions was multidimensional and changed over time, we could not use simple partitional clustering algorithms. Instead, we classified the participants’ time series by applying t-distributed stochastic neighbor embedding (t-SNE)[15] on the bottleneck layer output of autoencoder[16]. Then we chose uniform manifold approximation and projection (UMAP)[17] embedding focusing on two main components to find common patterns across all four affect dimensions. UMAP embedding results were plotted, and each group’s kernel density estimation plot was overlaid separately (Figs. 2e and g). The kernel density estimation plots replicated distinct patterns of each experienced emotion group that was in the original time series of the affect dimensions (see Supplementary Figs. 5 and 6). In other words, the spatial representations of stationary groups other than the RECe and RECx groups occupied distinct and non-overlapping regions, reflecting their consistent affect dimension values. In contrast, the representations of the RECe and RECx groups were more dispersed, with their regions spanning the entirety of the areas occupied by the stationary groups (Fig. 7b). This suggests that their responses are not confined to a specific range but rather traverse the entire spectrum from low to high as the fairness level increases. One-way ANOVA showed significant differences among the expected reward and reward acceptance as well as the four affect dimensions among the groups (p < 0.001, Figs. 2b, d, f, and h). Post-hoc TukeyHSD also showed significance (see Supplementary Fig. 7, Supplementary Table 2–11). Taken together, the participants’ expected and experienced emotions formed clusters that show both distinct temporal patterns and an aggregate summary of the affect dimension values.

### Reward and emotion resemble but are also distinct from each other

Both the original time series and embedding results exhibited parallel trajectories and an equivalent number of clusters for rewards and emotions. We further aimed to probe for unique variabilities within the rewards and emotions. Identifying independent fluctuations in the two variables can help establish whether the rewards and emotions are completely nested. To this end, we first plotted the estimated probability density function of the reward groups on the emotion UMAP embedding space shown in Fig. 2. The stationary reward groups occupied similar regions as the matched emotion groups, whereas the dynamic groups showed similar patterns of more dispersed spatial representations that spanned the entire range of the UMAP emotion space (Supplementary Fig. 8). The spatial representations of the reward groups were generally less confined than those of the emotion groups. In other words, the reward groups showed both alignment with and dispersion from the matched emotion groups on the latent space. Notably, the kernel density estimation was plotted using the same set of arguments as that used for the plot shown in Fig 2.

### Emotion PEs with prosocial dimensions as well as reward PE predict UG decisions

To replicate the result of [6] and to compare the predictive ability of our novel multi-dimensional dynamical ARM measure’s with that of the traditional affect grid model[6, 7, 12], a generalised linear mixed model (GLMM) was applied on the participants’ decision for each trial as dependent variable. Reward PE, valence PE, arousal PE, focus PE, and dominance PE were used as predictors with participants as the grouping variables. This design was adopted from previous work, and it allowed us to distinguish between the contributions of reward PE and emotion PEs during UG social interactions (Equation (1)). The models were nested based on the magnitude of Akiake Information Criteria (AIC) value reduction and compared using AIC for predictive accuracy[18]. The significance of model comparisons was tested via likelihood ratio tests (see Supplementary Tables 12–16).

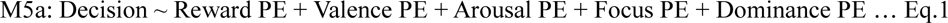

The results revealed that model M5a adopting all emotion PEs as well as reward PEs explained participant responses significantly better than the other models with simpler structures. Estimation on the beta coefficient of each predictor showed that the participants rejected more often when they received a smaller reward and felt less pleased and less in control than expected. A likelihood ratio test showed significant improvement in the explanatory power when the reward, dominance, focus, arousal PEs were sequentially added to the valence PE (χ^2^(4) = 495.83, P < 0.001, χ^2^(5) = 350.57, P < 0.001, χ^2^(6) = 131.75, P < 0.001, χ^2^(7) = 89.37, P < 0.001). The proportional Z-test for beta coefficients revealed that the valence PE outperformed all other PEs including the reward PE (β_intercept_ = 1.31, β_RPE_ = 0.60, β_VPE_ = 0.91, β_APE_ = -0.01, β_FPE_ = 0.11, β_DPE_ = 0.21). The dominance PE showed a relatively small contribution (Fig. 3; see Supplementary Table 17). This means that the participants adjusted their decision most significantly when they experienced greater unexpected (dis)pleasure, while their adjustments were less pronounced in response to the expectation violation of the reward and least pronounced in response to the unexpected level of dominance. Predictors of the model did not show high intra-individual-level correlations (see Supplementary Table 18), and their variance inflation factor (VIF) statistics indicated low collinearity with each other in the winning model (VIF_RPE_ = 1.04, VIF_VPE_ = 1.12, VIF_APE_ = 1.08, VIF_FPE_ = 1.21, VIF_DPE_ = 1.13). These results support the reliability of the relationship between the reward PE, emotion PEs, and the participants’ social decisions.

**Figure 3.**
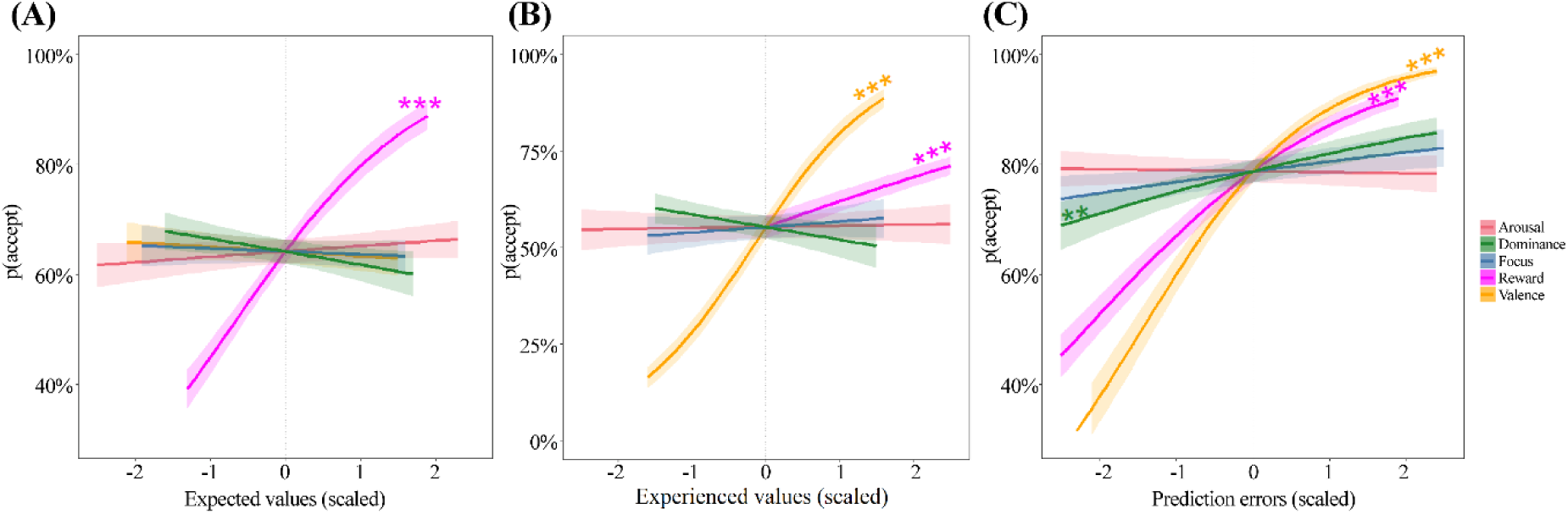
Relationship between respondent choice of acceptance and GLMM prediction. (A) Expected rewards and emotions as predictors. The expected reward predicts responder choice, as shown in Supplementary Table 19. (B) Offer and experienced emotions as predictors. The experienced valence and the offer from the proposer predict responder choice (Supplementary Table 20). (C) Reward and emotion PEs as predictors. The valence, reward, and dominance PEs contribute significantly to responder choice (Supplementary Table 17). ** p < 0.01, *** p < 0.001.

Applying GLMM separately for each experienced emotion group revealed that the groups reacted differently when the reward and emotions did not match their expectations (see Supplementary Table 22). The NON group rejected more often only when they felt less pleased than expected. The IND group showed more demand for prosociality by rejecting more when the reward and valence were smaller than their expectations. The RECx group’s reactions showed the greatest complexity as they rejected more frequently when they obtained a smaller reward and experienced less pleasure and less affective control than their anticipation. The RAT group exhibited greater disapproval only in response to a negative reward PE. The contributions of the focus PE and arousal PE to the social decision were not significant in any group. The predictive ability of the winning models was generally good, with all AUC values greater than 0.9 (Supplementary Fig. 9).

### Expected reward groups show different dynamics of mind-altering

Experienced emotion groups showed distinct modal trajectories of affect dimensions with changes in the fairness level (see Supplementary Figs. 6, and 10). To further test the group’s predictive ability in the dynamics of mind-altering, we compared the proportion of change from the expected reward groups to the experienced emotion groups (Fig. 4). Since both groups are semantically matched, this comparison corresponds to a change from an expectation about external rewards to the experienced emotions upon receiving actual rewards. The proportions of group change were 0.39 (PES), 0.46 (FAIR), 0 (RECe), and 0.13 (OPT). A chi-squared test revealed significant differences (p < 0.001), and the post-hoc z-tests revealed that all groups had p-values less than 0.05, except for the comparison between PES group and FAIR group (p = 0.159).

**Figure 4.**
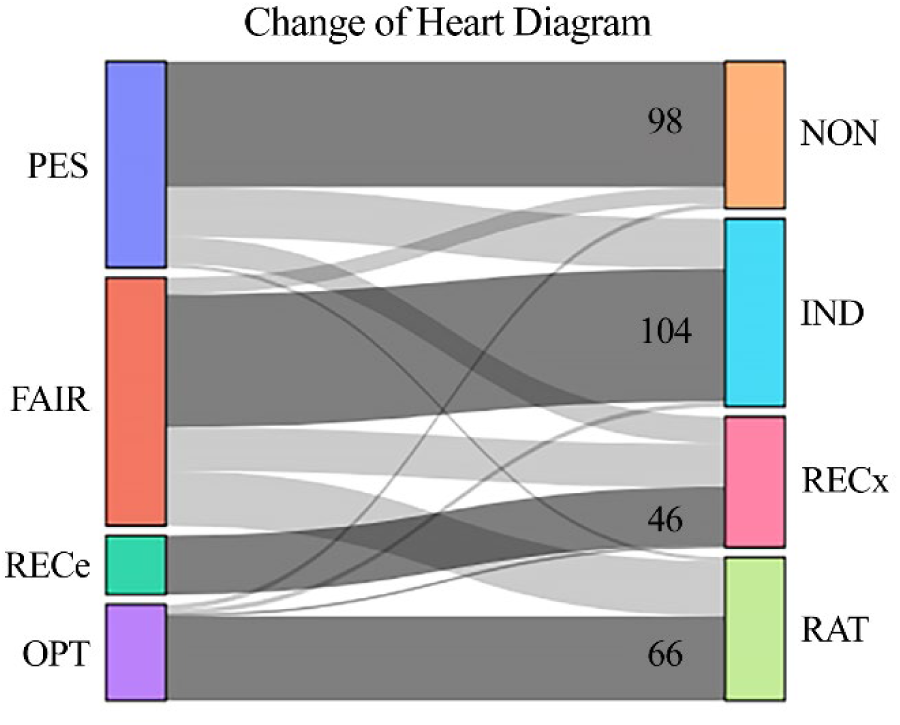
Sankey diagram of the change in membership from the expected reward group before the offer (left column) to the experienced emotion group upon receiving the offer (right column). Participants who are in the same group before and after the offer are represented by darker grey lines, with their numbers indicated. Abbreviations: PES, pessimistic expectation group; FAIR, fair expectation group; RECe, reciprocal expectation group; OPT, optimistic expectation group; NON, non-cooperative group; IND, indifferent group; RECx, reciprocal experience group; RAT, rational group.

In summary, the emotion clusters were identified through unsupervised classification based on the participants’ conscious appraisal of the basic–prosocial affect dimensions. These clusters exhibited distinct trajectories of basic-prosocial affect dimensions in relation to the participants’ experience of partner reciprocity that was embodied in the reward PE and fairness of offer. They also demonstrated differential variations in social decision-making that were accounted for by the predictive emotions along basic-prosocial affect dimensions. In other words, we found a comprehensive linkage between predictive emotions, social decision-making, and sense of reciprocity during UG.

### Focus and dominance dimensions show utility in differentiating and operationalising prosocial emotions

To properly evaluate the impact of focus and dominance dimensions on participant choices across different fairness levels, we conducted a detailed examination of their temporal patterns. First, we analysed the central tendencies of the affect dimensions for each experienced emotion group (see Supplementary Fig. 6). We found that within each cluster, the dimensions valence, focus, and dominance evolved in a similar fashion. However, when comparing across clusters, each cluster displayed distinct evolutionary trajectories for these dimensions. This suggests that although internal consistency exists within each cluster for valence, focus, and dominance, there is a clear divergence when comparing between different clusters. In contrast, these groups showed non-differentiable stationary central tendencies for the arousal dimension.

These central patterns assume that each dimension follows a predominant trend within the group. To substantiate this assumption, we graphed the probability energy distributions for each dimension (Fig. 5). We estimated the probability density function of affect dimensions and subsequently transformed these estimates using their negative logarithms. Notably, participants’ arousal values exhibited a pronounced valley in this transformed space. This indicates that the distribution of arousal is sharply peaked around its mode, suggesting a more constrained range of experiences (Fig. 5). This was true for all clusters. Moreover, distinct central patterns of valence, focus, and dominance for each group were replicated in the transformed space (see Supplementary Fig. 10).

**Figure 5.**
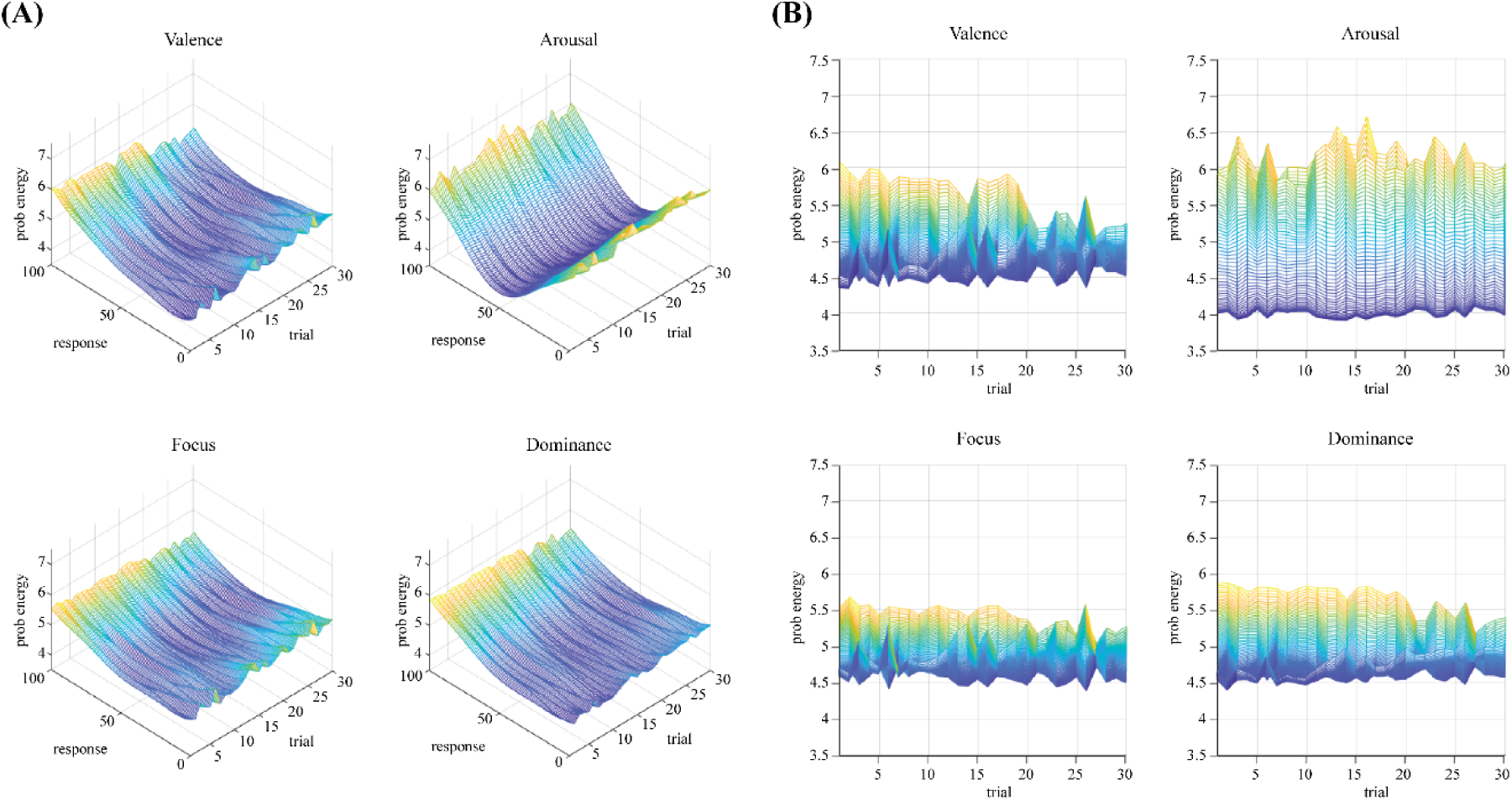
Probability energy distributions of the affective dimensions with increasing fairness levels. (A) Probability density function estimated based on the experienced emotions of all participants in each trial. The negative logarithm of the estimated probability is plotted, revealing a pronounced valley for the arousal dimension. (B) Each dimension was viewed directly along the response axis. The pronounced valley shown in (A) results in a larger height, indicating more densely distributed values centered around the mode.

From these results, the focus and dominance dimensions appear to have distinct modal trajectories within each cluster that can be isolated by our classification algorithm. We also plotted the energy landscape from the MSD distribution as in [19] (“egocentric approach”). The resultant energy landscapes were almost identical to each other, regardless of group labels and affect dimensions (see Supplementary Fig. 11).

To further evaluate the utility of focus and dominance in differentiating prosocial emotion labels, estimated probability density functions were plotted for the participants’ ratings of 9 prosocial emotions from ECT. We found supporting evidence for positive valence and self focus for pride, and negative valence and other focus for righteous anger, contempt, and moral disgust. However, no evidence was found supporting self-focus for shame, guilt, embarrassment, nor was other focus supported for elevation and gratitude (Fig. 6, see Supplementary Fig. 12). Prosocial emotions also occupied distinguishable areas along the affect grid from basic emotions (see Supplementary Fig. 13).

**Figure 6.**
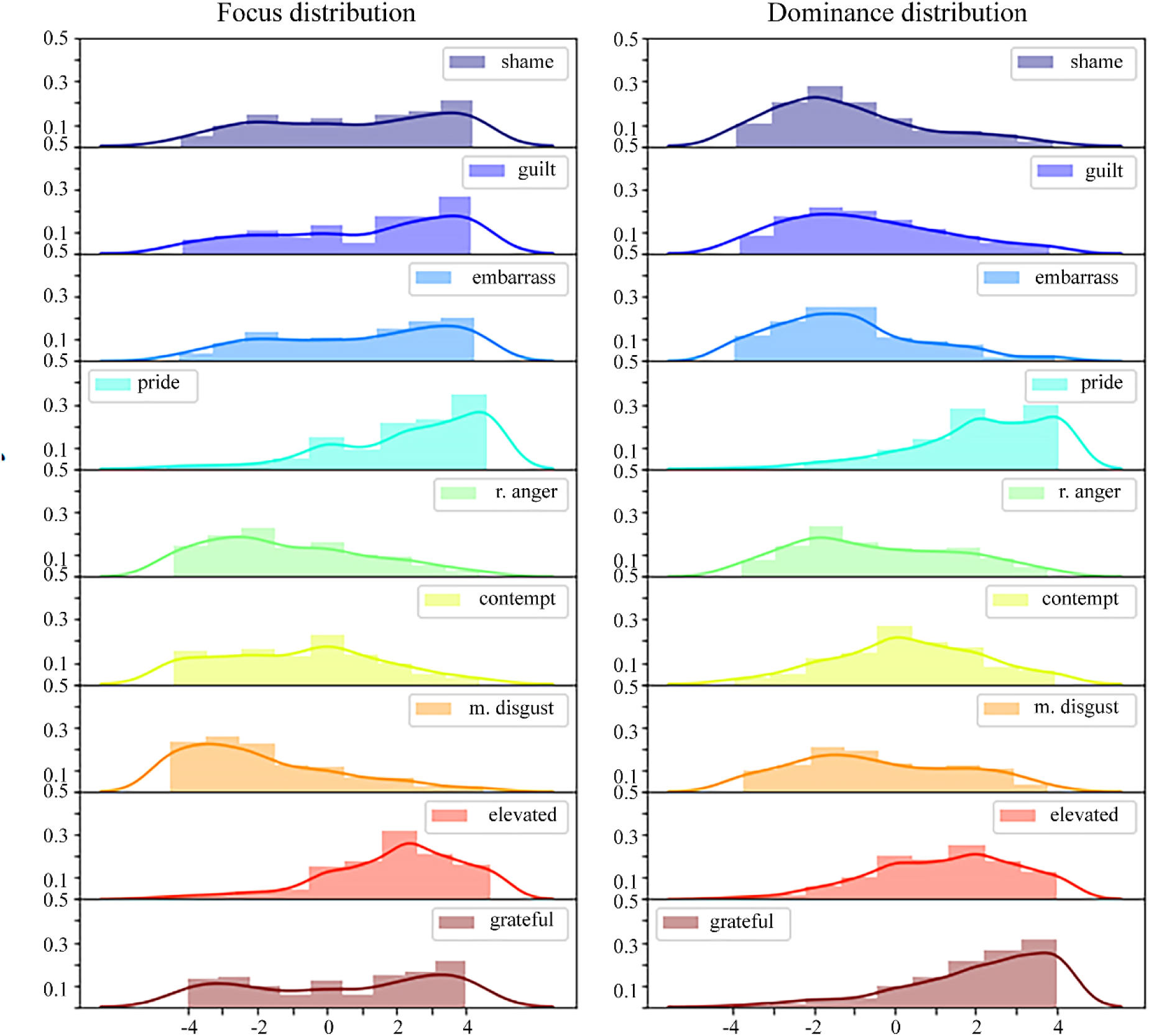
Kernel density estimation plots of focus and dominance for 9 moral emotions. The ratings of focus of the participants exhibit bimodal distributions for 3 negative self-focused emotions (shame, guilt, and embarrassment). Participants experienced self-focus for elevation and equivocal focus for gratitude. The Seaborn distplot function was used with the following arguments: hist = True, kde = True, rug = False, bins = 9.

## Discussion

We tested two main hypotheses:1) prediction error on prosocial affect dimensions, as well as prediction error on reward and basic affect dimensions, significantly contribute to a responder’s social decision during UG and that 2) the relationship between social decision-making, sense of reciprocity, and predictive emotion will differ across individuals. GLMM analyses on participant choices showed that adopting the focus and dominance dimension significantly improved the model’s explanatory power, and that dominance PE predicted participant choices. Additionally, the unsupervised classification algorithm identified inherent subsets of participants that exhibited distinct trajectories of reward and emotion experience that changed as a function of fairness level. These groups also differed in their relationships between reciprocity, basic-prosocial emotion, and social decision-making. We report for the first time that focus and dominance dimensions significantly contribute to both explaining participant social choices and identifying a subset of individuals from the UG task. Prosocial affect dimensions also showed utility in differentiating and operationalising prosocial emotions.

A major contribution of our study is that we were able to connect prediction errors along basic and prosocial affect dimensions to social decision-making, which was also linked to a sense of subjective (reward prediction error) and objective (fairness level) experience of partner’s reciprocity, using additional affect dimensions that capture aspects of social interaction. This indicates that conscious appraisal of basic-prosocial affects and their violations influence social decision-making, both in conjunction with and independently of sense of reciprocity.

Contemporary theories define emotion as an affective state of discrete reaction to internal or external events. Emotion changes as a function of situational meaning and its appraisal[20, 21] and it works as a component in subjective value (reward) calculation and driving action tendency[22]. Previous studies have shown that affective states including emotions [21, 23-25] and their anticipation [26-28] have a bidirectional causal relationship with decision-making [22, 29-31]. Specifically, social decision-making studies have adopted neuroeconomic tasks, and have reported a significant influence on affective states [3, 32-35]. Emotion influence on human behaviour was also exemplified by experimental studies using other [36-39] paradigms.

A relationship between reciprocity and social decision making has also been reported [1-3]. The subjective sense of reciprocity, which was embodied as reward prediction error, has been extensively studied for its contribution to associative learning [40-44]. This influence was replicated in the field of social decision making [4, 5]. However, the comprehensive linkage spanning all three phenomena of basic-prosocial emotion, social decision, and reciprocity remains poorly understood. The few studies that have ventured into this territory have only offered a limited perspective on the possible underlying dimensions related to social interaction that connect these phenomena [6, 12]. We introduce additional social affect dimensions that provide a more holistic understanding and, using a multi-methodological approach, further substantiate the intricate linkage binding these three phenomena.

In our study, prediction errors of valence and dominance, as well as of reward, had influence on social decisions, whereas arousal and focus prediction errors did not. This is in line with previous findings from [6], where valence had outweighed reward and arousal. However, unlike [6], arousal PE had no significant influence on participant choices. We believe our results support the interpretation of Heffner, Son, and FeldmanHall (2021) that emotion PE’s influence was context-dependent. In the previous study, participants at risk of depression exhibited a weakened relationship between emotion PE and punitive behaviours. Considering that our sample participants’ average PHQ-9 scores corresponded to a moderately severe level, the non-significant contribution of arousal PE is in line with previous reports (see Supplementary Table 1).

Acknowledging that the association between emotion PEs and participant choices may be influenced by specific contexts, we predicted that the relationship could differ among distinct populations. Thus, we applied GLMM to each experienced emotion group separately. Each group exhibited unique coefficient patterns (Supplementary Table 22). Non-cooperative group (NON) participants decided to reject offers more frequently when pleasure did not meet their expectation. Given a consistently low valence and other affect dimensions, the NON group may be in a state of negative, deactivated emotion, preventing them from adjusting interaction with partners until experiencing a relief of current displeasure. Moreover, reward PE had a non-significant contribution to responder choices. Hence, to change this group’s UG behaviour, an intervention should focus more on how pleased they feel rather than how much reward is optimal.

The rational group (RAT), on the other hand, increased punishment only for unexpectedly low offers. Participants in this group consistently reported high scores across all affect dimensions, potentially diminishing the marginal increases within these already positive scores. Furthermore, because the contribution of other responses was negligible, participants did not exhibit significant reactions to affect dimensions that fell below their expectations. This suggests that the group’s behaviour is influenced solely by reward PE without being swayed by their emotional experiences. The group seemed to consider only economic rationality during UG decision making, regardless of how they felt about their partner’s reciprocity. It is noteworthy that this group also showed the highest average scores in state anxiety, emotion regulation, and shame-proneness (see Supplementary Table 23). A possible explanation is that they more actively regulated their emotions and were more attuned to discrepancies between their self-image and moral standards, and these efforts could have contributed to their increased state anxiety. These cognitive efforts might be required to maintain their positive emotional state.

The indifferent group (IND) showed a more complex relationship. They rejected more often when they were either less pleased or less rewarded than expected. This relationship was also the weakest of all groups (see Supplementary table 22). Taken together with relatively neutral values of emotions during the task, this group could have been in a state of emotional detachment from their partners. The group with the most complex emotion trajectory (RECx group) also exhibited the most complex relationship between emotion PEs and social decision. They responded to unexpected levels of reward, pleasure, or affective control to adjust how they would respond to proposer offers. While their affect trajectories followed the variation of proposer’s offers, GLMM results suggest they changed their actions considering their own emotional state independently of reward PE.

These results suggest that each group can differentially react to monetary rewards and affective interventions. In other words, behavioural tasks can be individualised to influence participant’s social interaction in a disciplined way based on their emotional characteristics.

An additional dimension of focus was drawn from moral emotion study[8]. Although it failed to predict participant choices in any group, the models’ explanatory powers showed significant increases when adopting focus PE as the predictor for all groups. Focus also proved to be useful in differentiating moral emotions except for negative self-referential emotions of shame, guilt, and embarrassment.

The dominance dimension showed utility in predicting participant behaviours during UG as well as increasing models’ explanatory powers. It also seems promising for transdiagnostically phenotyping individuals, as only the RECx group reacted to dominance PE to change their behaviours. The use of the unsupervised classification algorithm implies that there are inheret subsets of participants[45] who can be isolated, and that these individuals exhibit distinct characteristics in their linkage between prosocial emotion, social decision, and sense of reciprocity. Definitions of affect dimensions and each emotion were informed to participants, ruling out the possibility of misinterpretation.

The higher-order theory of emotion posits that subjective experiences of emotions are inextricably linked to autobiographical information, encapsulated by the notion ‘no self, no emotion’[46]. Similarly, the theory of constructed emotions argues that emotions are not innate responses but socially generated constructs of affective experiences, which require knowledge of their source and the elements of control[47]. The explanatory power and predictive ability of dynamical ARM measure in predicting human behaviour increased by adopting social dimensions, which was evidenced by GLMM. This suggests that affective representation studies can indeed benefit from additional dimensions capturing autobiographical nature of emotion.

The study’s second significant contribution is that our classification algorithm can capture distinct temporal behaviours and aggregate summary of reward expectation, social decision, and expected and experienced emotions, suggesting its utility as a marker for experiential phenomena[46] as well as social decision making. Groups also exhibited differences in their experienced proportion of mind altering.

It is also noteworthy that participants were grouped into an equivalent number of classes with similar trajectories for expected reward, reward acceptance, and expected and experienced emotions. These similarities are inherent in the conscious appraisal of subjective experiences from independently and internationally recruited participants. UMAP embedding results also showed significant overlap between reward groups and emotion groups on latent space. In other words, similarity between temporal behaviours of subjective value calculation and emotion experiences carried over to latent space (see Supplementary Fig. 8). These similarities were also replicated using multiple dimensionality reduction algorithms (Fig. 7, see Supplementary Figs. 14 and 15)[15-17, 45, 48].

**Figure 7.**
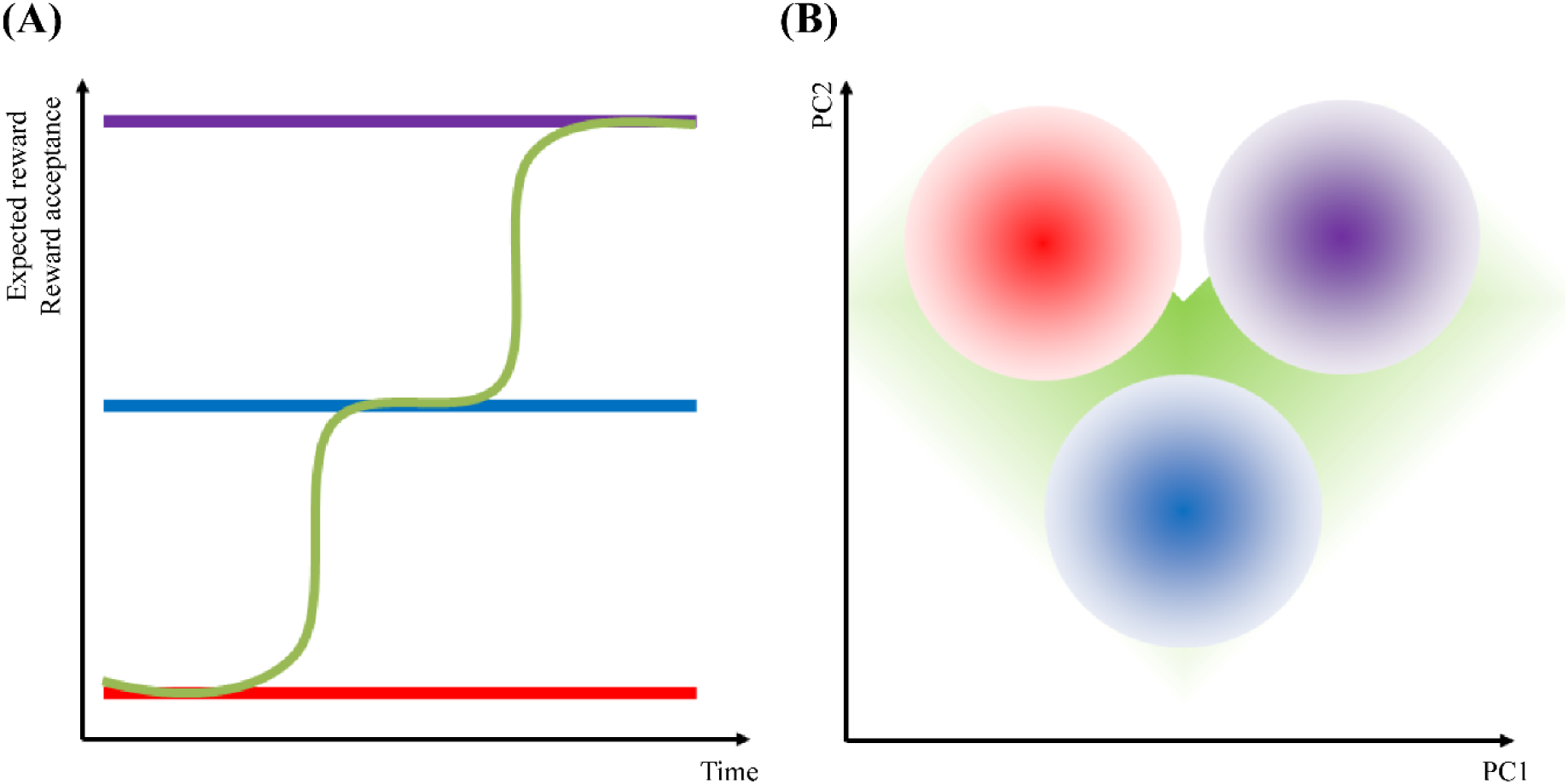
Conceptual diagram shows the similarity between the (A) temporal behaviours of the reward groups and the (B) spatial pattern of regions occupied by emotion groups in latent space. Groups of the same color across (A) and (B) exhibit similar trajectories but do not represent the same participants. The non-cooperator (red), indifferent (blue), and rational (purple) groups occupy distinct regions, whereas the reciprocal group (green) spans the regions of the other groups.

Additionally, reward PE and emotion PEs were independently predicting social decision making. Taken together, it seems that emotion is closely related to rewards but is not just internal proxy of external reward experience, corroborating the findings reported in [6]. A possible explanation is that emotion neural circuits, potentially integrating outputs from reward neural circuits along with signals related to autobiographical and interoceptive[49] information, may serve as a partial mediator by relaying these to decision-making neural circuits. This is analogous to the account of fear processing from the higher-order theory of emotion consciousness[46].

It is unclear whether these group labels represent the fluctuating affective states or more stable affective traits of individuals. Repeated measurements are needed to answer this question[50]. Because mental states are thought to be heterogeneous phenomena with a nested structure[51], dynamical ARM and our clustering method can be used for characterising other specific conditions such as psychiatric disorders.

Embedding a multi-dimensional time series into a lower dimensional state space offers comparability to the dynamics of subjective experiences, which is based on data science[45] and the network account of brain activity[19, 52]. The energy landscape showed differences in flatness of event rarity among different brain states in [19], which were analogous to activation energy changes in chemical reactions. It served as a neural marker of brain state dynamics; we have adopted this for a similar purpose for the state of subjective emotion experience. Despite the apparent differences in modal trajectories of affect dimensions, the energy landscape plots were nearly identical regardless of groups or affect dimensions. Mean squared displacement (MSD) measures the average response variation across different time lags and intervals. By analyzing the MSD value distribution, we assess the infrequency of these average response changes. The near-identical nature of the MSD plots across different groups, and across affect dimensions within groups, implies that the fundamental principles driving the emotion changes remain consistent across all groups, despite the distinct modal trajectories for emotional experience (see Supplementary Figs. 10, 11).

We were able to represent complex phenomena of moral emotion experiences using dimensions of valence, arousal, focus, and dominance (see Supplementary Fig. 12). The results showed a good agreement with the definition of some moral emotions, but not with shame, guilt, embarrassment (unclear self-focus), elevation, and gratitude (unclear other-focus) from [8] (see Supplementary Table 24). To the best of our knowledge, this is the first report on affective representation for moral emotions. Some moral emotions seem to need a redefinition. Further, shame, guilt, and embarrassment representations were almost identical to each other. Shame is defined as a negative evaluation of global self, guilt as a negative evaluation of specific actions, and embarrassment as a state of mortification following social predicaments[3, 53, 54]. These definitions support the cognitive emotion theory’s premise that emotions are functions of internal or external events, their situational meaning, and an agent’s appraisal. Prosocial emotions, being complex affective states, are significantly influenced by cognitive processes and extend beyond mere interoceptive or autobiographical signals[20, 21, 46, 47].

Our study has several limitations. First, verification of participants’ self-reported charactersitics and level of engagement was challenging due to online recruitment methods. Specifically, the Prolific platform collects demographic information prior to study participation. When participants initiated the experiment, the PsychoPy task opened in a new window, unlinked to their Prolific ID. Consent was only obtained upon completing the task, leading to a mismatch between the number of participants who shared demographic information and those who consented to participate in the study. Despite these issues, we advocate for more diverse samples to address the universality of emotional experiences and concerns about the heterogeneity problem in psychiatry[51], which we believe partially offsets the limitations of using an online platform. Second, a relatively high severity of psychopathology ratings was also noticeable. In fact, approximately 20% of the participants indicated that they had at least one on-going mental illness or mental health issue. Again, the universality of emotion experiences does not preclude populations with adverse mental health conditions, although stratified samples would be more desirable. Third, our task design uses a conscious recollection of emotion experience, leaving out behavioural and physiological responses from the analyses. It should be noted that all emotion groups experienced a narrow range of arousal centred around the zero point (Figs. 2f, h). We think that modifying the task to have more physiologically arousing inputs should be considered in future studies. Fourth, our main questions were not concerned with economic or organisational behavioural properties such as profit maximisation, communication, and conflict resolution[1-5]. Investigation into these aspects could provide more insights into human behaviours. Finally, momentary emotion labels can be predicted from social decision trajectories[12] to further predict prosocial behaviours.

In this study, we established a comprehensive linkage between basic-prosocial emotions, social decision making, and perceived reciprocity of a partner. We also proposed a novel data-driven classification algorithm for internal states of reward and emotion along with an affective representation method with additional variables drawn from the social psychology domain. We showed that this algorithm can identify groups with distinct trajectories of subjective experiences. Affect dimensions capturing aspects of social interaction seem to aid both the classification problem and prediction problem of human experience. It can also be utilised in testing and refining our knowledge of social constructs, which have only been subjectively defined. We hope our new methods can contribute to the fields of affect and value representation study, social psychology, and precision psychiatry.

## Methods

### Ethics Statement

This study was reviewed and approved by the Korea Advanced Institute of Science and Technology (KAIST) Institutional Review Board, Assurance # KH2021-084. Participant consent was collected explicitly upon task completion.

### Participants

A total of 636 participants were initially recruited through Prolific, an online psychological experiment platform (URL: https://www.prolific.co/). Among them, 89 returned before completing the emotion classification task and 51 did not complete the entire experiments, leaving 496 intact responses apart from some missing values. Further, 19 participants retracted their consent upon completing the task and one participant did not provide consent. The remaining 476 participants were considered for the subsequent analyses. In the prescreening phase of our research, some individuals provided consent for the use of their demographic data but opted not to participate in the main study. While their demographic information is included in the general dataset, no specific study-related data or outcomes from these individuals were collected or analysed. Demographic variables were obtained for all participants except for 113 who revoked their consent for providing demographic information (see Supplementary Table 27). Demographic information was obtained and de-identified in compliance with Prolific’s regulations. Nationalities were classified into regions according to [55]. Participants were internationally recruited, and were predominantly from Europe, Africa, and America regions. The nationalities were summarised for the top-5 countries and the rest were classified as Etc.

Clinical ratings were obtained at the end of experiment, and included Patient Health Questionnaire-9 (PHQ-9) for depression[56], Generalised Anxiety Disorder-7 (GAD-7) for anxiety[57], State-Trait Anxiety Inventory (STAI-X1) for state anxiety[58], Emotion Regulation Questionnaire (ERQ) for emotion regulation[59], and Personal Feelings Questionnaire-2 (PFQ-2) for shame and guilt[60] (see Supplementary Table 1). Missing values in clinical ratings were imputed with the average of each participant’s intact responses. The participants’ mean score of depression, anxiety, and state anxiety corresponded to moderately severe, moderate, and moderate level, respectively.

### Experimental procedures

The participants took part in an in-house computerised cognitive task, which consisted of three stages: ECT, dynamical ARM with ultimatum game (UG), and clinical ratings questionnaires. The task design referred to [6, 12] and utilised open-source PsychoPy programs, which were subsequently modified by the authors[61, 62] (Fig. 1). For the details of experimental procedures, see Supplementary Methods & Results.

### Emotion classification task

At the beginning of the experiment, the participants were asked to rate 18 different discrete emotions using valence, arousal, focus, and dominance to evaluate the utility of the affect dimensions in representing emotions (Fig. 1a)[7]. Nine basic emotions and nine moral emotions in adjective form were considered to be of interest. Eight of the basic emotions, except for fear, were drawn from the octant of emotion classification plots from [12] for enhanced separation. As five of the eight labels overlapped with six basic emotions[9], the authors included fear, resulting in nine basic emotions. Nine moral emotions were drawn from [8]. Missing values on the emotion classification task were excluded from the analyses using listwise deletion, leaving only 476 intact responses. The probability density function was estimated and plotted using Python’s matplotlib and seaborn packages (Fig. 6; see also Supplementary Figs. 12, 13).

Because of the input format’s potential influence on responses, valence and arousal were acquired using a traditional 2D affect grid with 501 * 501 resolution, which was delineated by a horizontal valence axis and a vertical arousal axis, while focus and dominance were acquired separately using individual 9-point Likert scales, ranging from -4 (completely toward others) to +4 (completely toward myself) for focus, and from -4 (completely out of control) to +4 (completely in control) for dominance. The definitions of each affect dimension and emotion label were explained to the participants at the beginning of experiment.

### Dynamical affective representation mapping with ultimatum game

The participants went through 30 trials of the one-shot ultimatum game as a responder (Fig. 1b). The trials were divided into three blocks of 10 trials each with different levels of fairness: unfair (mean offer 0.9), moderate (mean offer 2.8), and fair (mean offer 4.9) blocks with range of offers from $0–7. The rewards in each block were pseudo-randomised values with a standard deviation of 1.0. The blocks were presented in a completely random order. The participants were told that they would interact with a new person in each trial and that the blocks represented distinct groups with unique human-like figures. Earned credits were automatically accumulated in dollar amounts and were given to participants in cents with the numerical value being the same as earned credits. A predictive mean matching algorithm was used to impute the missing values in rewards and affect dimensions using R’s mice() package. Analyses were done after realigning the randomised blocks in an ascending order of fairness.

At the beginning of each trial, the participants were asked to respond with expected rewards in a 0 ∼ 9 range, and the affect dimension they would feel if given the expected amount (Fig. 1b). Then, the pseudo-randomised offers were given to the participants. After being presented with the offer, the participants were asked to rate their current affect dimensions, following which they decided either to accept or reject the offer. The effect of fairness order on overall acceptance proportion (p_accept_) was tested with six sets of randomised block order as a predictor using OLS regression (Equation (2)) with non-significant differences.

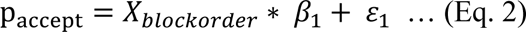

### Clinical ratings questionnaires

Upon completing dynamical ARM with UG, the participants rated their severity of depression (PHQ-9), anxiety (GAD-), state anxiety (STAI-X-1), emotion regulation (ERQ), and shame and guilt (PFQ-2) (see Supplementary Table 1). The possible effect of clinical ratings on social decision-making was also evaluated. First, we divided the participants based on the lower and upper quartiles of each rating. Then, independent t-tests were applied to compare p_accept_ along all levels of offer for two groups. None of the 40 comparisons were significant (see Supplementary Table 25). Earned credits were also compared, and only ERQ score groups showed trend-level significant differences (p = 0.084).

### Unsupervised clustering algorithms reveal common data structure across time series of reward and emotions

K-means clustering was applied to the entire time series of participants’ expected reward and reward acceptance, rather than individual time points. Based on inertia and silhouette score, K = 4 was selected (see Supplementary Figs. 1, 2). To identify groups of distinct emotion trajectories with fairness level changes, t-SNE was applied to the autoencoder’s bottleneck output using TensorFlow [15, 16, 63]. Supplementary Fig. 3 visualises the t-SNE embedding with two components from the autoencoder’s hidden layer output. Similar to reward, four clusters were identified from inertia and silhouette scores (see Supplementary Figs. 4–6). The mean values of affect dimensions and number of participants in each group are summarised in Supplementary Table 26. To compare the average expected reward, reward acceptance, and expected and experienced emotions between groups, one-way ANOVA and post-hoc TukeyHSD were applied.

Dimensionality reduction algorithms (PCA, t-SNE[15], MDS[45], UMAP[17], tPHATE[48]) were applied to the time series of expected and experienced emotions using scikit-learn, TensorFlow, Keras, umap, and tphate packages (Fig. 2, also see Supplementary Figs. 14, 15). The kernel density estimation plot was visualised separately for each emotion group using Python’s matplotlib and seaborn packages.

### Reward and emotion resemble but are also distinguished from each other

Reward prediction error (RPE) and emotion prediction error (EPE) were standardised but not mean-centered as zero prediction error implied an accurate prediction[6]. RPEs’ and EPEs’ contributions to social decision were investigated by a generalised linear mixed model (GLMM) using the glmer function from R’s lme4 package. Because our study involved an exploratory analysis, the Akaike information criterion (AIC) was used for model comparison[18]. A log likelihood test on deviances was performed for significance of model comparison. The beta coefficients for each predictor in the winning models were evaluated to test the significance of contributions from each prediction error terms (see Supplementary Table 22).

### Focus and dominance dimensions show utility in differentiating and operationalising prosocial emotions

Using the egocentric approach from [19], we developed the energy landscape on response change probability distribution using mean squared displacement. Moreover, the probability energy distribution on state space of trial-by-response was generated. This approach allows an investigation of both dynamics of transition probability of emotion independent from environmental constraints, and dynamics of context-specific emotion (Fig. 5, see also Supplementary Figs. 10, 11).

### Statistical Analyses

All statistical analyses were performed using Python, R, and MATLAB. Participant responses were handled using the pandas library and NumPy package. Matplotlib and seaborn packages were used for visualisation and kernel density estimation. Ordinary least squares estimation and linear mixed model were applied via statsmodels and scipy packages. One-way ANOVA and independent t-test or TukeyHSD were used to compare the differences in mean values using stats and scipy. The time series clustering algorithm and corresponding packages were as follows: kmeans, kmedoids, PCA, tSNE, MDS (scikit-learn), Autoencoder (TensorFlow, Keras), UMAP (umap), and tPHATE (tphate). A predictive mean matching algorithm was used to impute the missing values in rewards and affect dimensions using the mice() package in R. GLMM was performed using the glmer function in R’s lme4 library, and the energy landscape analyses used a MATLAB code that was modified from [19].

## Acknowledgements

This study was supported by the Medical Scientist Training Program from the Ministry of Science & ICT of Korea, the Brain Research Program through the National Research Foundation of Korea (NRF) funded by the Ministry of Science & ICT (NRF-2022M3E5E8081200)

## Author contributions

J.K. and B.J. conceived the original idea, planned and designed the experiments. J.K. performed the experiments, derived the models and analysed the data. All authors contributed to designing the research, the interpretation of the results. J.K. and B.J. wrote the paper with input from all authors. B.J. supervised the project.

## Competing interests

The authors declare no competing interests.

## Additional Information

**Supplementary information** The online version will contain supplementary material available upon publication of this manuscript.

**Correspondence and requests for materials** should be addressed to Bumseok Jeong.

## Data availability

The minimum dataset supporting the findings of this study is currently available to peer reviewers upon request during the review process. The dataset will be made publicly accessible in a GitHub repository upon publication.

## Notes

### Competing Interest Statement

The authors have declared no competing interest.

